# FGF19 and its analog Aldafermin cooperate with MYC to induce aggressive hepatocarcinogenesis

**DOI:** 10.1101/2023.09.15.557921

**Authors:** José Ursic-Bedoya, Guillaume Desandré, Carine Chavey, Pauline Marie, Arnaud Polizzi, Benjamin Rivière, Hervé Guillou, Eric Assenat, Urszula Hibner, Damien Gregoire

## Abstract

FGF19 hormone has pleiotropic metabolic functions, notably the modulation of insulin sensitivity, glucose/lipid metabolism and energy homeostasis. On top of its physiological metabolic role, FGF19 has been identified as a potentially targetable oncogenic driver, notably in hepatocellular carcinoma (HCC). Nevertheless, FGF19 remained an attractive candidate for treatment of metabolic disease, prompting the development of analogs uncoupling its metabolic and tumor-promoting activities.

Using pre-clinical mice models of somatic mutation driven HCC, we assessed the oncogenicity of FGF19 in combination with frequent HCC tumorigenic alterations: p53 inactivation, CTNNB1 mutation, CCND1 or MYC overexpression. Our data revealed a strong oncogenic cooperation between FGF19 and MYC. Most importantly, we show that this oncogenic synergy is conserved with a FGF19-analog Aldafermin (NGM282), designed to solely mimic the hormone’s metabolic functions. In particular, even a short systemic treatment with recombinant proteins triggered rapid appearance of proliferative foci of MYC-expressing hepatocytes.

The fact that FGF19 analog Aldafermin is not fully devoid of the hormone’s oncogenic properties raises concerns in the context of its potential use for patients with damaged, mutation-prone liver.

## Introduction

Fibroblast Growth Factor 19 (FGF19) is an ileum-secreted hormone that controls bile acids synthesis and regulates several metabolic functions (Owen *et al*, 2015). Pleiotropic activities of FGF19, and its rodent ortholog FGF15, are important regulators of liver physiology, through increase of metabolic rate, improvement of glucose and insulin tolerance, and decrease of fasting insulin levels (Lan *et al*, 2017; Marcelin *et al*, 2014; Gadaleta & Moschetta, 2019). On the other hand, FGF19 displays a mitogenic activity and has been identified as an oncogenic driver in hepatocellular carcinoma (HCC). Focal amplification of 11q13 locus, encompassing *FGF19*, is among the most frequent amplification events in HCC (6-15% of HCC) (Sawey *et al*, 2011; Schulze *et al*, 2015; Guichard *et al*, 2012). This genetic alteration is associated with more aggressive tumors, higher risk of recurrence after surgical resection and lower overall survival rates (Miura *et al*, 2012; Ahn *et al*, 2014; Schulze *et al*, 2015; Kang *et al*, 2019). Moreover, mice with forced FGF19 overexpression develop hepatic tumors (Nicholes *et al*, 2002; Zhou *et al*, 2014a, 2017c) while targeting of FGFR4, the main FGF19 receptor in the liver, reduces tumor growth (Hagel *et al*, 2015; French *et al*, 2012). Importantly, an inhibitor of FGFR4, Fisogatinib (BLU-554), has demonstrated clinical activity in HCC patients with aberrant FGF19 expression (Kim *et al*, 2019) (phase I clinical trial, NCT02508467).

The interconnection between FGF19 oncogenic and metabolic activities, as well as the relative contributions of the downstream effectors of the hormone, remain poorly characterized. FGF19 binding to its receptor FGFR4 and co-receptor β-Klotho (KLB) has been reported to activate several major signal transduction pathways, such as Ras/MAPK, Akt, Mst1/2 and β-catenin signaling, which are all implicated in hepatic carcinogenesis (Chen *et al*, 2019; Ornitz & Itoh, 2015; Ji *et al*, 2019). Moreover, FGF19 acts on multiple cell types and binds other members of the FGFR family, including FGFR1c (Owen *et al*, 2015; Lan *et al*, 2017). This redundancy between FGFRs may account for some limitations in the efficacy of FGFR4 specific inhibitors (Tao et al, 2022).

Despite the clear association between FGF19 and hepatocellular carcinoma, the metabolic properties of the hormone, notably its remarkable inhibitory effect on lipid accumulation, have prompted efforts to develop analogues devoid of the oncogenic activity. (Gadaleta *et al*, 2018; Zhou *et al*, 2014b). NGM282 (more recently named Aldafermin), has been reported to retain bile acid synthesis inhibition, without promoting tumor formation (Zhou *et al*, 2014b, 2017a; DePaoli *et al*, 2019). Aldafermin, which was selected in a systematic screen of FGF19 mutants in diabetic *db/db* mice, carries a 5-amino acid deletion and 3 amino acid substitutions in the N-terminal part of the protein, which is the FGFR4 receptor binding domain (Zhou *et al*, 2014b). The proposed interpretation for the reported loss of the oncogenic activity of Aldafermin was that the altered binding to the FGFR4/KLB receptor would fail to activate the STAT3 pathway, which might be implicated in the oncogenic properties of FGF19 (Zhou *et al*, 2017c, 2014b). Aldafermin was subsequently tested in several clinical trials involving patients receiving the molecule subcutaneously during 12 or 24 weeks for nonalcoholic steatohepatitis (NASH) without cirrhosis, primary sclerosing cholangitis, primary biliary cholangitis or bile acid diarrhea (Hirschfield *et al*, 2019; Harrison *et al*, 2022; BouSaba *et al*, 2023; Mayo *et al*, 2018). These first trials generated mixed results. A phase 2b randomized clinical trial involving 160 patients with NASH compensated cirrhosis treated with Aldafermin for 48 weeks is ongoing (NCT04210245).

Here, we investigated the effects of FGF19/15 and Aldafermin in the framework of oncogenic cooperation with several common HCC drivers. We reasoned that a diseased, inflamed liver is subjected to microenvironmental insults that may result in increased somatic mutation incidence, as shown in the context of cirrhosis (Rebouissou & Nault, 2020). It was therefore of interest to assess possible tumor promoting effects of oncogenic combinations involving FGF19 and its analogs.

## Results

### FGF19 and FGF15 cooperate with MYC to trigger hepatic carcinogenesis

We used hydrodynamic gene transfer (HGT) (Zhang *et al*, 1999; Liu *et al*, 1999) to transfect hepatocytes *in vivo* to combine overexpression of the human FGF19 with genetic alterations frequently observed in HCC. We selected *Trp53* inactivation, β-catenin mutation and MYC overexpression (Zucman-Rossi *et al*, 2015) (Figure 1A), as well as the overexpression of cyclin D1 (CCND1), the latter reproducing the focal amplification of 11q13 observed in 10% of HCC (Sawey *et al*, 2011).

**Figure 1:**
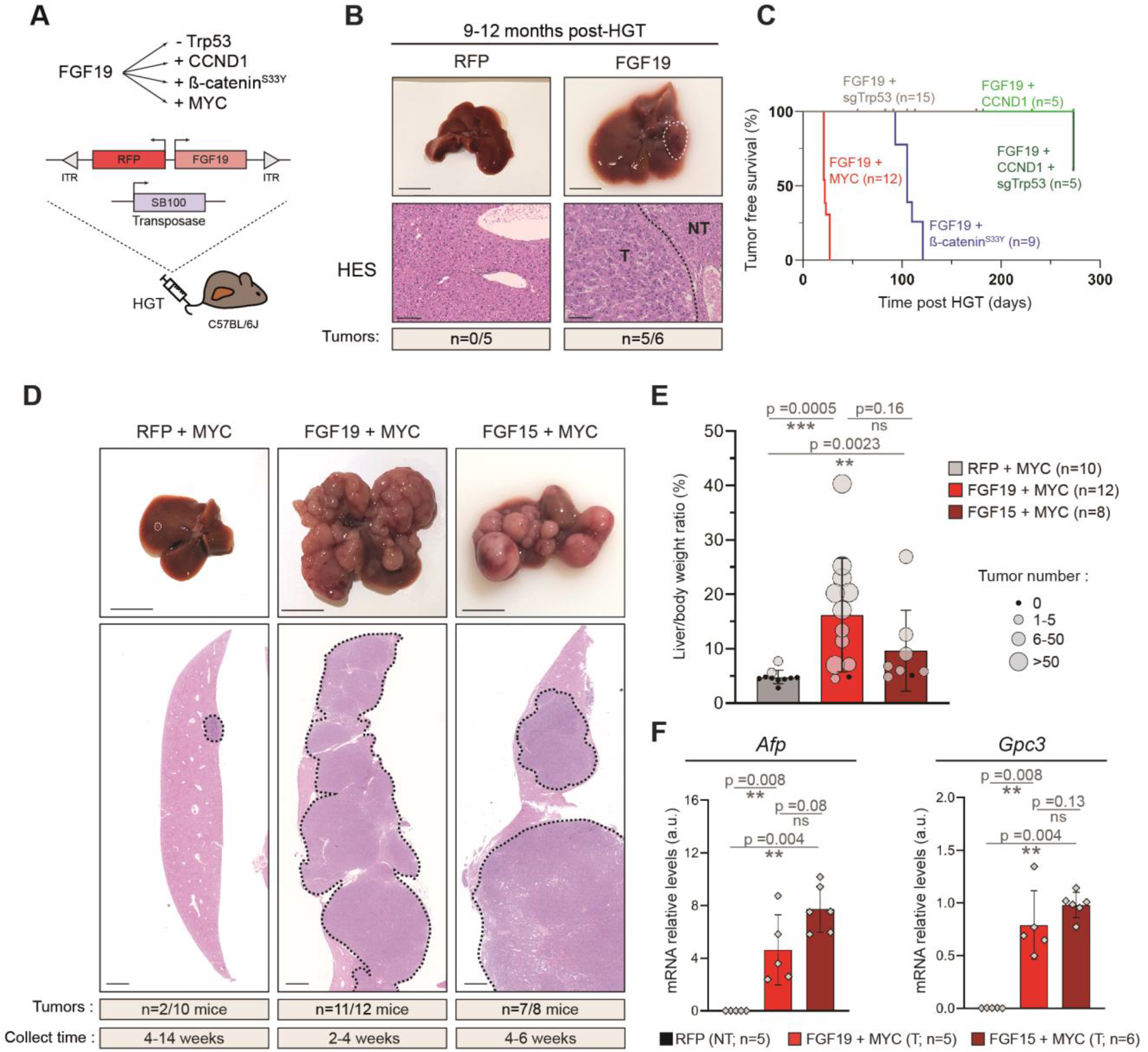
FGF19 and FGF15 cooperate with MYC to trigger liver carcinogenesis. **A**. Mouse model strategy based on hydrodynamic gene transfer (HGT) with Sleeping-Beauty transposase 100 (SB100). **B**. Representative liver and HES staining after HGT with empty vector or FGF19. Scale-bar: 1cm (liver picture), 100 μm (HES section). **C**. Tumor-free survival curve (Kaplan-Meier method) for FGF19+MYC, FGF19+sgTrp53, FGF19+CCND1, FGF19+CCND1+sgTrp53, FGF19+ß-cateninS33Y groups. **D**. Representative liver and HES staining after HGT with MYC+RFP, FGF19+MYC and FGF15+MYC. Scale-bar: 1cm. **E**. Liver/body weight ratio and tumor number per liver in mice injected with MYC + RFP, FGF19 + MYC and FGF15 + MYC. **F**. Gene expression analyses by RT-qPCR of hepatocellular carcinoma markers Glypican 3 and Alpha fetoprotein in RFP parenchyma, FGF19+MYC and FGF15+ MYC tumors. Mann-Whitney test statistical significance is indicated. Abbreviations: HES: Hematoxylin & Eosin & Saffron. RFP: red fluorescent protein. CCND1: cyclin D1. sgTrp53: single-guide Tumor-related protein 53. T: tumor. NT: non-tumor parenchyma. Gpc3: glypican 3. Afp: alpha fetoprotein.

FGF19 transfection of hepatocytes led to supra-physiological plasma levels of the hormone (1-10 ng/mL), that persisted over several months and displayed long-term metabolic effects(Ursic-Bedoya *et al*, 2022). Nine to twelve-months after HGT, most animals developed tumors (incidence n=5/6, one tumor/liver) (Figure 1B). Thus, with circulating FGF19 levels 6 to 100-fold lower than in previously described mouse models of FGF19 overexpression (Zhou *et al*, 2017b; Nicholes *et al*, 2002), we confirmed the previously reported moderate oncogenic effect of FGF19.

Associating FGF19 overexpression with Crispr/Cas9-mediated inactivation of p53 (Bacevic *et al*, 2019), did not accelerate tumorigenesis (Figure 1C). Similarly, the incidence of tumor development in FGF19/CCND1/sg*Trp53* animals ten months after HGT remained identical to that of FGF19 animals (n=2/5, maximum of 1 tumor/liver) (Figure 1C). Therefore, our results suggest that there is no oncogenic cooperation between FGF19 overexpression and p53 inactivation or CCND1 overexpression.

In contrast, combination of overexpression of FGF19 and β-catenin^S33Y^ gave rise to tumors in 4 months (n=9/9, mean= 3.7 tumors/liver, Figure 1C), significantly increasing hepatic carcinogenesis that was triggered by β-cateninS33Y alone at the same time point (n=2/6, 1 or 2 tumors/liver).

However, the most spectacular results were obtained when FGF19 was associated with the overexpression of MYC: at a maximum of four weeks after HGT, these animals had to be sacrificed, for ethical reasons (Figure 1C). Their livers presented an elevated tumor burden (n=11/12; mean= 72 tumors per liver, CI_95_ [45, 99]) (Figure 1D, E). This was in sharp contrast with control mice transfected with MYC and RFP (i.e., control empty vector) for which only 2 mice out of 10 presented tumors (1 tumor per liver), despite the extended delay of their sacrifice, up to 14 weeks post-HGT (Figure 1D, E).

FGF15, the rodent ortholog of FGF19 was reported to be devoid of oncogenic properties, presumably due to structural differences with the human hormone (Zhou *et al*, 2017b). However, the co-expression of FGF15 with MYC also gave rise to hepatic carcinogenesis, albeit within a slightly longer timeframe (7/8 mice exhibited tumors after 4-6 weeks post-HGT) and a lower tumor burden (mean= 10.3) in comparison with FGF19+MYC (Figure 1D, E). Anatomopathological analyses revealed that the FGF19+MYC and FGF15+MYC tumors were both moderately differentiated and these tumors expressed hepatocellular carcinoma markers Alpha-fetoprotein (*Afp*) and Glypican 3 (*Gpc3*) (Figure 1F). We conclude that FGF19 and FGF15 cooperate with MYC to trigger rapid and aggressive hepatic carcinogenesis.

### Aldafermin is oncogenic when combined with MYC overexpression

Our discovery of significant oncogenic activity of FGF15 prompted us to have a closer look at FGF19 analog Aldafermin (NGM282). Of note, the sequence homology between FGF19 and Aldafermin is high, considerably higher than between the human and mouse orthologs. Indeed, there are only 8 AA differences between the natural hormone and its pharmacological analog and the AlphaFold predicted 3D structures of the two proteins display 97% similarity (Figure 2A)(Jumper *et al*, 2021).

**Figure 2:**
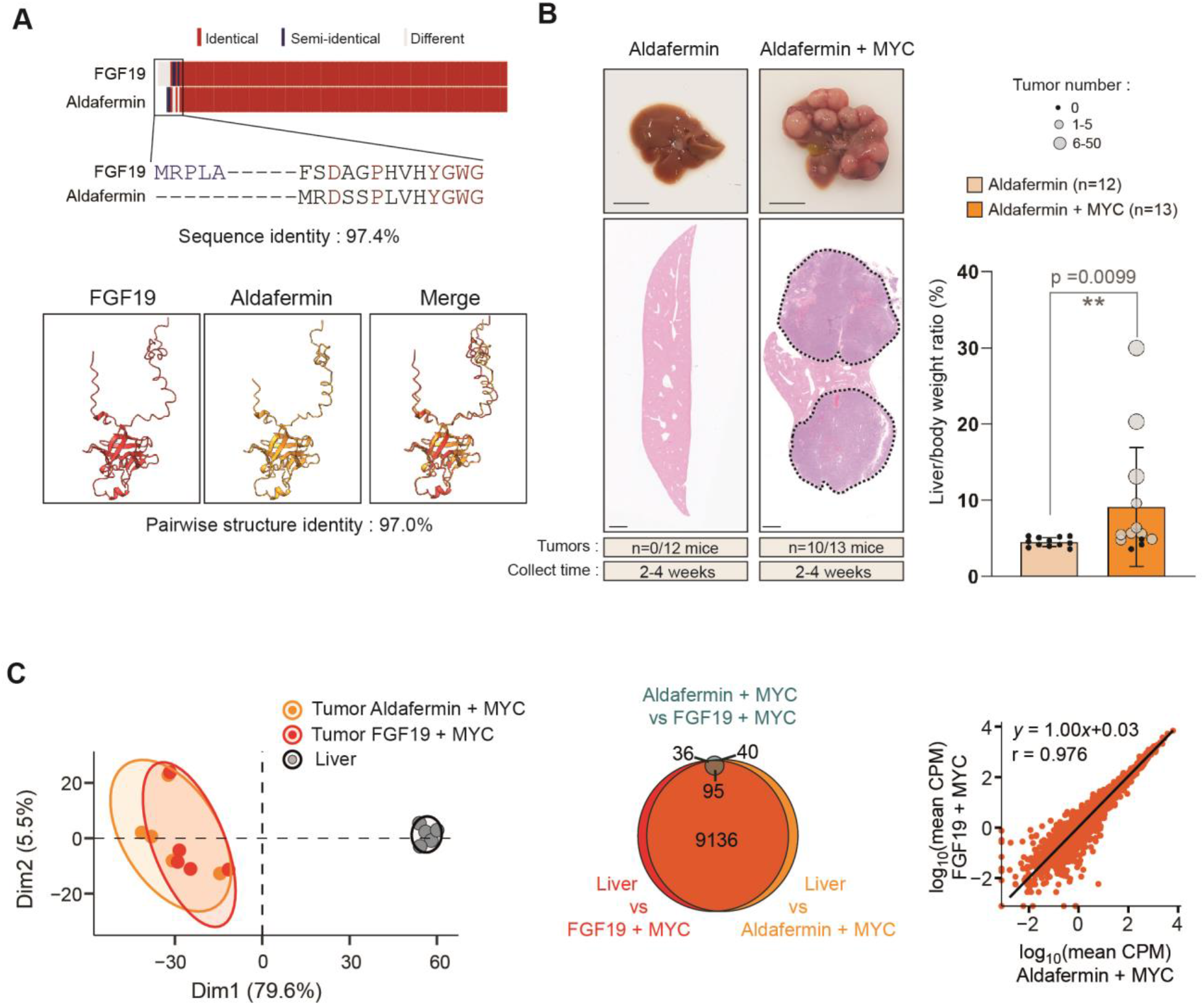
Aldafermin and FGF19 are indistinguishable in cooperating with MYC to promote liver carcinogenesis. **A**. Upper panel: sequence alignment of FGF19 and Aldafermin proteins. Lower panel: 3D AlphaFold prediction of proteins showing almost identical structure and folding. **B**. Left panel: Representative image of liver and HES staining in liver sections after HGT with Aldafermin or Aldafermin+MYC, 2 to 4 weeks after transfection. Scale-bar: 1cm. Right panel: Liver/body weight ratio and tumor number per liver in mice injected with Aldafermin or Aldafermin+MYC. **C**. Principal Component Analysis of RNAseq gene expression profiles of tumors obtained after HGT with FGF19+MYC or Aldafermin+MYC or liver as control. Euler diagram of the differential analysis results from RNAseq. Each circle represents the number of genes with an adjusted pvalue<0.05 for each comparison. Correlation plot of the mean log10(CPM) of the Aldafermin+MYC and FGF19 +MYC tumors. Each point represents one gene, and the black line is a linear regression over the whole dataset. Equation of the linear regression and its Pearson’s correlation coefficient are indicated. Mann-Whitney test statistical significance is indicated. Abbreviations: HES: Hematoxylin & Eosin & Saffron.

We therefore tested the oncogenicity of Aldafermin hepatic expression, either alone or in combination with MYC (Figure 2B). As expected, Aldafermin overexpression alone did not trigger tumor development after 2 to 4 weeks (n=0/12). In striking contrast, the majority of livers co-transfected with Aldafermin and MYC exhibited multiple tumors between 2 and 4 weeks post-HGT (n=10/13; tumors mean number=7.9; CI_95_ [1.3,14.5]). HES staining revealed that tumors were moderately differentiated and were in fact indistinguishable from the FGF15/19 tumors by an anatomopathological examination (Figure 2B). To further examine this point, we performed gene expression analyses of the tumors by bulk-RNAseq (n=5 for each). Principal Component Analysis of gene expression profiles, Euler diagram and correlation analysis illustrate that Aldafermin+MYC and FGF19+MYC tumors indistinguishable transcriptomic profiles (Figure 2C).

Overall, our results reveal that the modification of 8 AA at the N-terminal part of FGF19, producing the FGF19-analog Aldafermin, does not abolish oncogenic properties of the hormone, at least in the context of co-expression with MYC.

### Oncogenic effects of systemic administration of FGF19 and Aldafermin recombinant proteins

While FGF19 is expressed by hepatocytes in a significant proportion of HCC, hepatic expression of a transgene encoding Aldafermin is clearly not physiological. We therefore designed an experimental scheme to test the mitogenic effects of Aldafermin and FGF19 in a setup more closely resembling the clinical reality, where patients receive Aldafermin daily (Harrison *et al*, 2020; Mayo *et al*, 2018; Hirschfield *et al*, 2019; Harrison *et al*, 2018).

We produced tag-free recombinant FGF19 and Aldafermin (Choi *et al*, 2023, 2020) (Figure 3A) and performed intra-peritoneal injections of 2μg of purified proteins per mouse (corresponding to the 3 mg of the drug given to patients in clinical trials(Harrison *et al*, 2018)). Transcriptional repression of hepatic *Cyp7a1*, a *bona fide* target of FGF19, confirmed the biological activity of the recombinant proteins *in vivo* (Figure 3B). A single injection led to circulating levels of around 20ng/mL after two hours, in accordance with the reported short half-life of the hormone(Degirolamo *et al*, 2016). To approach plasma levels observed in Aldafermin-treated patients, we next injected 1.2μg of recombinant proteins every 12 hours. We tested the effect of FGF19 or Aldafermin administration on animals previously subjected to hepatocyte transfection by HGT with MYC plasmid (Figure 3C).

**Figure 3:**
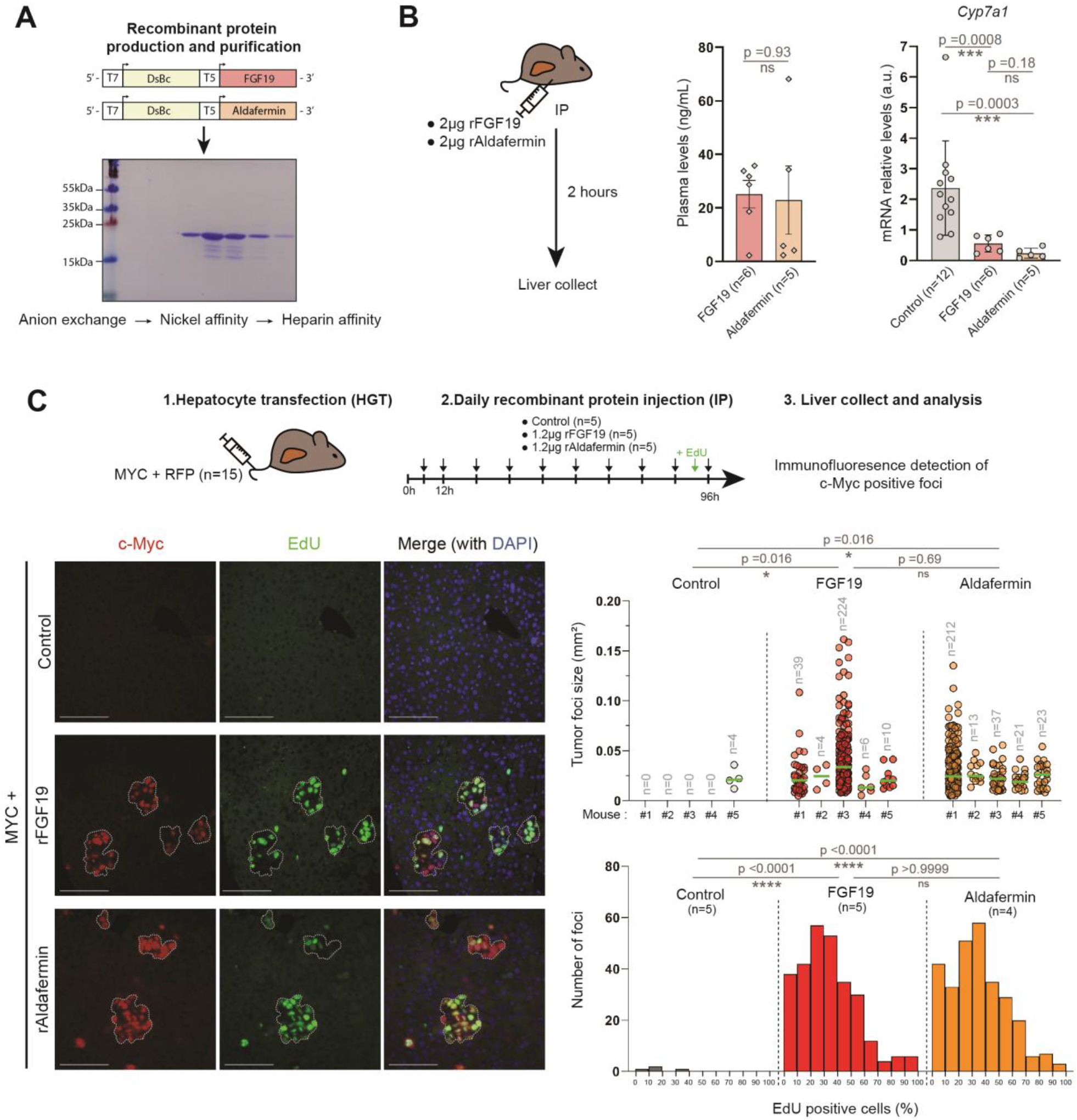
Systemic administration of recombinant FGF19 and Aldafermin recapitulates cooperation with MYC. **A**. Schematic representation of the workflow for recombinant FGF19 and Aldafermin production and purification. **B**. Plasma levels of FGF19/Aldafermin and Cyp7a1 relative mRNA level 2 hours after intraperitoneal injection of 2μg of recombinant FGF19 or Aldafermin. **C**. Upper panel: Schematic representation of the experiment. After in vivo transfection, mice were injected every 12 hours with either 1.2μg rFGF19, rAldafermin or PBS. Left panel: Immunofluorescence and Click-iT EdU assay performed on liver sections. Detoured are tumoral foci. Scale bar = 100μm. Right upper panel: Size of the MYC-positive foci detected in the analyzed liver slices. Green bar represents the mean of the values. Number of detected foci is indicated. Right lower panel: number of foci composed of 0 to 100% of EdU positive cells. Kolmogorov-Smirnov test statistical significance is indicated. Mann-Whitney test statistical significance is indicated. Abbreviations: DsBc: disulfide bond C, rFGF19: recombinant FGF19, rAldafermin: recombinant Aldafermin.

After 4 days, livers of mice from the control group (HGT-MYC followed by PBS injection) had no detectable tumoral foci and very few MYC positive cells, which was expected, since these cells are rapidly eliminated in the absence of a cooperating oncogenic event. In contrast, mice that received either FGF19 or Aldafermin presented numerous MYC-positive foci (mean size= 0.026mm^2^; CI_95_ [0.015, 0.037] and 0.026mm^2^; CI_95_ [0.020, 0.031] mm^2^ for FGF19 and Aldafermin respectively) (Figure 3C). EdU incorporation indicated that these foci were highly proliferative, and that Aldafermin effects were indistinguishable from FGF19 (Figure 3C). Thus, Aldafermin and FGF19 show similar pro-tumorigenic activities on MYC expressing hepatocytes, confirming that the metabolic and oncogenic activities of the analog of the hormone have not been fully uncoupled.

## Discussion

In this study, we have identified a strong oncogenic cooperation between the FGF19 hormone and MYC, driving hepatocarcinogenesis. Importantly, we discovered that FGF19 analog Aldafermin conserves oncogenic properties in this context.

Among all the combinations tested, FGF19 and MYC oncogenic cooperation stood out, as it triggered an impressively fast and aggressive tumorigenesis. As FGF19 tumor-promoting effects were observed in the context of cooperation with MYC or β-catenin, both of which have a documented pro-mitogenic activity, our result suggests that they are unlikely to be solely based on stimulation of cell proliferation. The precise molecular mechanisms responsible for these synergistic effects remain to be elucidated. Of note, while aberrant FGF19 expression is frequent in HCC, other types of tumors, such as head and neck squamous cell carcinoma, have been shown to be driven by FGF19 (Gao *et al*, 2019)and our findings might also be of interest in these contexts.

Aldafermin is not the only FGF19 analog described as non-mitogenic. Other variants of the hormone that have been engineered (Wu *et al*, 2010; Gadaleta *et al*, 2018; DePaoli *et al*, 2019) all carry changes in the N-terminal part of the protein, as these modifications have been described to curtail the oncogenic activity of FGF19 due to a weakened activation (dimerization) of the FGFR4 receptor (Jin *et al*, 2022). However, our data indicate that, in the context of MYC overexpression, the signaling by Aldafermin to its downstream effectors is sufficient to promote oncogenic growth. Of note, considering the functional redundancy between FGF19 and FGF21, and their shared specificity for the FGFR receptors (FGFR4, FGFR1c), it may be of interest to further investigate possible oncogenic cooperation of FGF21 analogs.

Finally, it is important to consider the relevance of oncogenic cooperation in a clinical setting. Aldafermin, as well as other FGF19 analogs, are designed to treat patients with damaged liver. There is evidence for clonal expansion in NASH livers (Wang et al, 2023) and cirrhosis is associated with accumulation of mutations (Schulze *et al*, 2015). MYC target gene signature has recently been reported to be associated with non-cirrhotic NASH-induced HCC (Pinyol *et al*, 2021). Altogether, MYC amplification is found in 6-17% of human HCC (Ally *et al*, 2017), while MYC pathway is activated in close to 30% of human HCC (Kaposi-Novak *et al*, 2009), making it a relevant alteration in the context of oncogenic cooperation.

We believe that our results showing that Aldafermin retains oncogenic properties raise significant concerns. They indicate that the benefits of treating metabolic liver disorders with FGF19 analogs need to be carefully balanced against the drugs’ possible direct effects on carcinogenesis.

## Material and Methods

### Mice experiments

All reported animal procedures were carried out in accordance with the rules of the French Institutional Animal Care and Use Committee and European Community Council (2010/63/EU). Animal studies were approved by institutional ethical committee (Comité d’éthique en expérimentation animale Languedoc-Roussillon (#36)) and by the Ministère de l’Enseignement Supérieur, de la Recherche et de l’Innovation (D. Gregoire: APAFIS #32384-2021070917596346 v2). ARRIVE guidelines were followed.

### Hydrodynamic gene transfer

Hydrodynamic injections were performed in 6 to 8 weeks-old female mice as described previously (Zhang *et al*, 1999; Liu *et al*, 1999). Briefly, 0.1 mL/g of a solution of sterile saline (0.9% NaCl) containing plasmids of interest were injected into lateral tail vein in 8-10 s. 12.5 μg of LentiCRISPRv2-sg*Trp53*, pSBbi-RN-FGF19, pT3-EF1a-MYC, pSBBi-BB-CCND1, or pSBBi-RN-CTNNB1^S33Y^ were injected together with sleeping beauty transposase SB100X (ratio of 5:1). Livers were harvested when mice exhibited signs of tumor development or at a predetermined endpoint ranging from 4 to 52 weeks after injection.

### Plasmids

All plasmid sequences were validated by whole-plasmide sequencing (SNPsaurus). pCMV(CAT)T7-SB100 was a gift from Zsuzsanna Izsvak (Addgene plasmid #34879). SgTp53 sequence (from Zhang Lab database): 5’-ATAAGCCTGAAAATGTCTCC-3’, was cloned into LentiCRISPRv2 vector. Plasmids constructs pSBbi-RN-FGF19 and pSBbi-BB-FGF15 were respectively generated by cloning the human *FGF19* amplified from Huh7 cDNA and the mice *Fgf15* amplified from ileum cDNA onto the pSBbi-RN (Addgene #60519) and pSBbi-BB (Addgene #60521) plasmids digested by the SfiI restriction enzyme. Murine CCND1 and human CTNNB1^S33Y^ open reading frames were cloned into pSBbi-BB and pSBbi-RN, respectively, using the same strategy. Aldafermin (NGM282) coding sequence was generated from pSBbi-RN-FGF19 plasmid using site directed mutagenesis (Q5® Site-Directed Mutagenesis – New England BioLabs) using the primers FOR 5’-ATCGGGCCTCTGAGGCCATGCGCGACTCGTCGCCCCTCGTGCACTACGGCTGG-3’ and REV 5’ – CGATGGCCTGACAGGCCTTACTTCTCAAAGCTGGGACTCC– 3’.,

The plasmids for the bacterial expression of FGF19 and Aldafermin were generated by producing a sequence composed of the T5 promoter followed by the sequence coding for the disulfide bond isomerase (ΔssDsbC), then a T7 promoter followed by a codon-optimized variant of either FGF19 or Aldafermin CDS based on published work by Choi et al (Choi *et al*, 2023). Those sequences were subsequently cloned in the pQLinkG2 (Addgene #13671) using the XhoI/PasI restriction sites.

### Purification of recombinant proteins

Bacterial expression plasmids for rFGF19/rAldafermin were transformed into Rosetta-gami™ 2 (DE3) competent cells (Novagen). Bacteria were cultured in LB medium + 100μg/mL ampicillin and induced with 0.2mM IPTG at 30°C for 3h.

Recombinant proteins were purified without tags using methods described by Choi et al. (Choi et al, 2023, 2020). To summarize, dried pellets of induced bacteria were solubilized in 30mL of buffer A1 (20mM Tris-HCL pH8.0, 1mM DTT) then sonicated. Lysate was cleared by centrifugation and charged on a 5mL HiTrap Q HP column (Cytiva) then eluted using a continuous gradient of buffer B1 (20mM Tris-HCL pH8.0, 1mM DTT, 1M NaCl). Fractions containing the protein of interest were pooled,diluted 1:3 with buffer A2 (20mM Sodium phosphate pH7.0, 150mM NaCl pH7.0, 1mM imidazole) and charged on a 1mL HisTrap (Nickel) column. Protein was eluted with an 80mM imidazole concentration using a step-wise elution with buffer B2 (20mM Sodium phosphate pH7.0, 150mM NaCl pH7.0, 100mM imidazole). Fractions were pooled,diluted 1:6 with buffer A3 (20mM sodium phosphate pH6.5) and loaded on a 1mL HiTrap Heparin HP affinity column (Cytiva). Protein was eluted at a 600mM NaCl using stepwise gradient of buffer B3 (Buffer B: 20mM sodium phosphate, 1M NaCl, pH6.5). Protein purity was assayed using Coomassie blue, and protein quantification was assayed using human FGF19 ELISA kit (Biovendor, RD191107200R). Protein was then diluted 1:2 in PBS+0.2%BSA and stored at - 20°C.

### Histology assays, image acquisition and analysis

Livers were fixed for 24 h in 10% neutral buffered formalin, dehydrated, and embedded in paraffin and, cut into 3-μm-thick sections. Tissue sections were stained with hematoxylin, eosin saffron (HES) with Leica autostainer for preliminary analysis. Tumor differentiation and characteristics were reviewed by an expert pathologist (BR). Slides were digitally processed using the Nanozoomer scanner (Hamamatsu).

Immunofluorescence assays were performed after deparaffination and antigen retrieval in citrate buffer pH6.0 (Sigma Aldrich). The Click-iT AF488 (ThermoFisher Scientific) EdU detection reaction was then performed according to the manufacturer’s instructions. Labelling of c-Myc positive cells was performed with human c-Myc antibody (Abcam rabbit ab#32072 dilution 1:200) and anti-rabbit secondary antibody (Thermofisher anti-rabbit IgG AF633). Images were acquired using the Thunder Imager Tissue (Leica) microscope. Quantitative image analysis was performed using the QuPath v0.4.4 software. Foci were hand drawn and positive cell detection were performed using nuclear staining as basis of cell recognition.

### RNA isolation, qPCR and RNA-Seq analysis

The RNA was extracted from liver tissue and purified using RNeasy mini kit (Qiagen) according to manufacturer’s protocol. Reverse transcription of total RNA (1μg) was done with QuantiTect Reverse Transcription kit (Qiagen), and cDNA quantified using LC Fast start DNA Master SYBR Green I Mix (Roche) with primers detailed below on LightCycler480 apparatus (Roche). Gene expression levels were normalized with hypoxanthine phospho-ribosyltransferase (Hprt). Primer pairs used for qPCR:

Hprt_For 5’-GCAGTACAGCCCCAAAATGG-3’

Hprt_Rev 5’-GGTCCTTTTCACCAGCAAGCT-3’

Afp_For 5’- CTGTCTCAGTCATTCTAAGAATTGCT-3’

Afp_Rev 5’- CTCCTCGATGTGTTTCTGC-3’

Gpc3_For 5’-AGGACTGTGGCCGTATG-3’

Gpc3_Rev 5’-GCAATAACCACCGCAAGG-3’

Cyp7a1_For 5’-CTGCAACCTTCTGGAGCTTA-3’

Cyp7a1_Rev 5’- ATCTAGTACTGGCAGGTTGTTT-3’

For RNA-Seq analysis, RNA was extracted from snap frozen individual tumors, using RNeasy mini kit (Qiagen) with DNAse treatment. RNA integrity was validated using RNA BioAnalyzer (Agilent), all RIN > 9.0. The preparation of the library was done with the TruSeq Stranded mRNA Sample Preparation kit (Illumina). The sequencing was performed in an Illumina Hiseq 2500 sequencer by the Sequencing Platform of Montpellier (GenomiX, MGX, France; www.mgx.cnrs.fr), with 150 base pairs (bp) paired end reads to an estimated depth of 70million reads per sample. In order to perform a quality control of the sequencing, FastQC over the fastq files containing the raw reads. All the reads that passed the quality control were aligned to the mouse reference genome (Mus musculus NCBI Mm39) with HiSAT2 and the counts per gene were quantified using the tool htseq-count. The NCBI RefSeq mouse genome annotations (accessed 27/02/2023) were used for establishing the coordinates of each gene and their corresponding transcripts. Differential gene expression analysis was performed in R using the EdgeR package (Robinson et al, 2010).

Transcripts with less than one count per million (CPM) were discarded for the analysis. Fitting the globalized linear model (GLM) was done using the tagwise dispersion. Transcript with at least two comparisons under the adjusted p.value threshold of 0.05 were considered as differentially expressed genes (DEG).

### Protein modelling and alignment

Protein modelling of FGF19 and Aldafermin were done using AlphaFold predictions via the ChimeraX software (Pettersen et al, 2021). Protein 3D alignment was made using the RCSB PDB Pairwise Structure Alignment website (https://www.rcsb.org/alignment last access 20 July 2023) using the jFATCAT method (Ye & Godzik, 2003).

### Experimental design and Statistical Analysis

The investigators were blinded every time it was possible. Data sets were tested with 2-tailed unpaired Student t tests or Mann-Whitney U tests, correlations were analyzed with Pearson’s χ2 test using Prism Software version 8 (GraphPad). Significant P values are shown as: *P <0.05, **P <0.01, ***P <0.001, and ****P <0.0001.

Data visualization and statistical analysis were performed using R software version 3.5.1 (R Foundation for Statistical Computing, Vienna, Austria. https://www.R-project.org) and Bioconductor packages. Comparisons of the mRNA expression levels between groups were assessed using Mann-Whitney U test. Spearman’s rank-order correlation was used to test the association between continuous variables. Univariate survival analysis was performed using Kaplan-Meier curve with log-rank test.

## Acknowledgments

We acknowledge Montpellier Biocampus facilities: the imaging facility (MRI), the “Réseau d’Histologie Expérimentale de Montpellier” - RHEM facility supported by SIRIC Montpellier Cancer Grant INCa_Inserm_DGOS_12553, the european regional development foundation and the Occitanian region (FEDER-FSE 2014-2020 Languedoc Roussillon) for processing some of our animal tissues. We acknowledge the MGX facility for the RNAseq analysis, and the MRI imaging platform. We are grateful to Olivier Coux and Aymeric Bailly for their help in producing FGF19 and Aldafermin recombinant proteins, and to Geun-Joong Kim’s Lab for advice. We thank Georges-Philippe Pageaux for helpful discussions and Michael Hahne for support. This work was funded by EVA-Plan cancer INSERM THE, Association Française pour l’Etude du Foie (AFEF) AAP_2021 and supported by SIRIC Montpellier Cancer Grant INCa_Inserm_DGOS_12553, Fondation ARC and INCa (2019-134). JUB was funded by SIRIC Montpellier. The funders had no role in study design, data collection and analysis or publication process.

## Data availability

The RNA-sequencing data have been deposited in the Gene Expression Omnibus (GEO, NCBI) repository, and are accessible through GEO Series accession number GSE242953. All newly created materials are made available to the community.

## References

Ahn SM, Jang SJ, Shim JH, Kim D, Hong SM, Sung CO, Baek D, Haq F, Ansari AA, Lee SY, et al (2014) Genomic portrait of resectable hepatocellular carcinomas: Implications of RB1 and FGF19 aberrations for patient stratification. Hepatology 60: 1972–1982

Ally A, Balasundaram M, Carlsen R, Chuah E, Clarke A, Dhalla N, Holt RA, Jones SJM, Lee D, Ma Y, et al (2017) Comprehensive and Integrative Genomic Characterization of Hepatocellular Carcinoma. Cell 169: 1327–1341.e23

Bacevic K, Prieto S, Caruso S, Camasses A, Dubra G, Ursic-Bedoya J, Lozano A, Butterworth J, Zucman-Rossi J, Hibner U, et al (2019) CDK8 and CDK19 kinases have non-redundant oncogenic functions in hepatocellular carcinoma. bioRxiv

Chen J, Du F, Dang Y, Li X, Qian M, Feng W, Qiao C, Fan D, Nie Y, Wu K, et al (2019) Fibroblast Growth Factor 19– Mediated Up-regulation of SYR-Related High-Mobility Group Box 18 Promotes Hepatocellular Carcinoma Metastasis by Transactivating Fibroblast Growth Factor Receptor 4 and Fms-Related Tyrosine Kinase 4. Hepatology

Choi HJ, Cheong DE, Yoo SK & Kim GJ (2023) One-step metal affinity purification of recombinant hFGF19 without using tags. Protein Expr Purif 201

Choi HJ, Cheong DE, Yoo SK, Park J, Lee DH & Kim GJ (2020) Soluble expression of hfgf19 without fusion protein through synonymous codon substitutions and dsbc co-expression in e. Coli. Microorganisms 8: 1–12

Degirolamo C, Sabbà C & Moschetta A (2016) Therapeutic potential of the endocrine fibroblast growth factors FGF19, FGF21 and FGF23. Nat Rev Drug Discov 15: 51–69

DePaoli AM, Zhou M, Kaplan DD, Hunt SC, Adams TD, Marc Learned R, Tian H & Ling L (2019) FGF19 analog as a surgical factor mimetic that contributes to metabolic effects beyond glucose homeostasis. Diabetes 68: 1315–1328

French DM, Lin BC, Wang M, Adams C, Shek T, Hötzel K, Bolon B, Ferrando R, Blackmore C, Schroeder K, et al (2012) Targeting FGFR4 inhibits hepatocellular carcinoma in preclinical mouse models. PLoS One 7: 1–12

Gadaleta RM & Moschetta A (2019) Metabolic Messengers: fibroblast growth factor 15/19. Nat Metab 1: 588–594

Gadaleta RM, Scialpi N, Peres C, Cariello M, Ko B, Luo J, Porru E, Roda A, Sabbà C & Moschetta A (2018) Suppression of Hepatic Bile Acid Synthesis by a non-tumorigenic FGF19 analogue Protects Mice from Fibrosis and Hepatocarcinogenesis. Sci Rep 8: 1–10

Gao L, Lang L, Zhao X, Shay C, Shull AY & Teng Y (2019) FGF19 amplification reveals an oncogenic dependency upon autocrine FGF19/FGFR4 signaling in head and neck squamous cell carcinoma. Oncogene 38: 2394–2404

Guichard C, Amaddeo G, Imbeaud S, Ladeiro Y, Pelletier L, Maad I Ben, Calderaro J, Bioulac-Sage P, Letexier M, Degos F, et al (2012) Integrated analysis of somatic mutations and focal copy-number changes identifies key genes and pathways in hepatocellular carcinoma. Nat Genet 44: 694–698

Hagel M, Miduturu C, Sheets M, Rubin N, Weng W, Stransky N, Bifulco N, Kim JL, Hodous B, Brooijmans N, et al (2015) First selective small molecule inhibitor of FGFR4 for the treatment of hepatocellular carcinomas with an activated FGFR4 signaling pathway. Cancer Discov 5: 424–437

Harrison SA, Rinella ME, Abdelmalek MF, Trotter JF, Paredes AH, Arnold HL, Kugelmas M, Bashir MR, Jaros MJ, Ling L, et al (2018) NGM282 for treatment of non-alcoholic steatohepatitis: a multicentre, randomised, double-blind, placebocontrolled, phase 2 trial. The Lancet 391: 1174–1185

Harrison SA, Rossi SJ, Paredes AH, Trotter JF, Bashir MR, Guy CD, Banerjee R, Jaros MJ, Owers S, Baxter BA, et al (2020) NGM282 Improves Liver Fibrosis and Histology in 12 Weeks in Patients With Nonalcoholic Steatohepatitis. Hepatology 71: 1198–1212

Hirschfield GM, Chazouillères O, Drenth JP, Thorburn D, Harrison SA, Landis CS, Mayo MJ, Muir AJ, Trotter JF, Leeming DJ, et al (2019) Effect of NGM282, an FGF19 analogue, in primary sclerosing cholangitis: A multicenter, randomized, double-blind, placebo-controlled phase II trial. J Hepatol 70: 483–493

Ji S, Liu Q, Zhang S, Chen Q, Wang C, Zhang W, Xiao C, Li Y, Nian C, Li J, et al (2019) FGF15 Activates Hippo Signaling to Suppress Bile Acid Metabolism and Liver Tumorigenesis. Dev Cell 48: 460–474.e9

Jin L, Yang R, Geng L & Xu A (2022) Annual Review of Pharmacology and Toxicology Fibroblast Growth Factor-Based Pharmacotherapies for the Treatment of Obesity-Related Metabolic Complications.

Jumper J, Evans R, Pritzel A, Green T, Figurnov M, Ronneberger O, Tunyasuvunakool K, Bates R, Žídek A, Potapenko A, et al (2021) Highly accurate protein structure prediction with AlphaFold. Nature 596: 583–589

Kang HJ, Haq F, Sung CO, Choi J, Hong SM, Eo SH, Jeong HJ, Shin J, Shim JH, Lee HC, et al (2019) Characterization of hepatocellular carcinoma patients with FGF19 amplification assessed by fluorescence in situ hybridization: A large cohort study. Liver Cancer 8: 12–23

Kaposi-Novak P, Libbrecht L, Woo HG, Lee YH, Sears NC, Conner EA, Factor VM, Roskams T & Thorgeirsson SS (2009) Central role of c-Myc during malignant conversion in human hepatocarcinogenesis. Cancer Res 69: 2775–2782

Kim RD, Sarker D, Meyer T, Yau T, Macarulla T, Park JW, Choo SP, Hollebecque A, Sung MW, Lim HY, et al (2019) Firstin-human phase i study of fisogatinib (BLU-554) validates aberrant FGF19 signaling as a driver event in hepatocellular carcinoma. Cancer Discov 9: 1696–1707

Lan T, Morgan DA, Rahmouni K, Sonoda J, Fu X, Burgess SC, Holland WL, Kliewer SA & Mangelsdorf DJ (2017) FGF19, FGF21, and an FGFR1/β-Klotho-Activating Antibody Act on the Nervous System to Regulate Body Weight and Glycemia. Cell Metab 26: 709–718.e3

Liu F, Song Y & Liu D (1999) Hydrodynamics-based transfection in animals by systemic administration of plasmid DNA. Gene Ther 6: 1258–66

Marcelin G, Jo YH, Li X, Schwartz GJ, Zhang Y, Dun NJ, Lyu RM, Blouet C, Chang JK & Chua S (2014) Central action of FGF19 reduces hypothalamic AGRP/NPY neuron activity and improves glucose metabolism. Mol Metab 3: 19–28

Mayo MJ, Wigg AJ, Leggett BA, Arnold H, Thompson AJ, Weltman M, Carey EJ, Muir AJ, Ling L, Rossi SJ, et al (2018) NGM 282 for Treatment of Patients With Primary Biliary Cholangitis: A Multicenter, Randomized, Double-Blind, Placebo-Controlled Trial. Hepatol Commun 2: 1037–1050

Miura S, Mitsuhashi N, Shimizu H, Kimura F, Yoshidome H, Otsuka M, Kato A, Shida T, Okamura D & Miyazaki M (2012) Fibroblast growth factor 19 expression correlates with tumor progression and poorer prognosis of hepatocellular carcinoma. BMC Cancer 12

Nicholes K, Guillet S, Tomlinson E, Hillan K, Wright B, Frantz GD, Pham TA, Dillard-Telm L, Tsai SP, Stephan JP, et al (2002) A mouse model of hepatocellular carcinoma: Ectopic expression of fibroblast growth factor 19 in skeletal muscle of transgenic mice. American Journal of Pathology 160: 2295–2307

Ornitz DM & Itoh N (2015) The fibroblast growth factor signaling pathway. Wiley Interdiscip Rev Dev Biol 4: 215–266

Owen BM, Mangelsdorf DJ & Kliewer SA (2015) Tissue-specific actions of the metabolic hormones FGF15/19 and FGF21. Trends in Endocrinology and Metabolism 26: 22–29 doi:10.1016/j.tem.2014.10.002 [PREPRINT]

Pettersen EF, Goddard TD, Huang CC, Meng EC, Couch GS, Croll TI, Morris JH & Ferrin TE (2021) UCSF ChimeraX: Structure visualization for researchers, educators, and developers. Protein Science 30: 70–82

Pinyol R, Torrecilla S, Wang H, Montironi C, Piqué-Gili M, Torres-Martin M, Wei-Qiang L, Willoughby CE, Ramadori P, Andreu-Oller C, et al (2021) Molecular characterisation of hepatocellular carcinoma in patients with non-alcoholic steatohepatitis. J Hepatol 75: 865–878

Rebouissou S & Nault JC (2020) Advances in molecular classification and precision oncology in hepatocellular carcinoma. J Hepatol 72: 215–229

Robinson MD, McCarthy DJ & Smyth GK (2010) edgeR: a Bioconductor package for differential expression analysis of digital gene expression data. Bioinformatics 26: 139–40

Sawey ET, Chanrion M, Cai C, Wu G, Zhang J, Zender L, Zhao A, Busuttil RW, Yee H, Stein L, et al (2011) Identification of a therapeutic strategy targeting amplified FGF19 Supplement. Cancer Cell 19: 347–58

Schulze K, Imbeaud S, Letouzé E, Alexandrov LB, Calderaro J, Rebouissou S, Couchy G, Meiller C, Shinde J, Soysouvanh F, et al (2015) Exome sequencing of hepatocellular carcinomas identifies new mutational signatures and potential therapeutic targets. Nat Genet 47: 505–511

Tao Z, Cui Y, Xu X & Han T (2022) FGFR redundancy limits the efficacy of FGFR4-selective inhibitors in hepatocellular carcinoma.

Ursic-Bedoya J, Chavey C, Desandré G, Meunier L, Dupuy A-M, Gonzalez-Dopeso Reyes I, Tordjmann T, Assénat E, Hibner U & Gregoire D (2022) Fibroblast growth factor 19 stimulates water intake. Mol Metab: 60

Wang Z, Zhu S, Jia Y, Wang Y, Kubota N, Fujiwara N, Gordillo R, Lewis C, Zhu M, Sharma T, et al (2023) Positive selection of somatically mutated clones identifies adaptive pathways in metabolic liver disease. Cell 186: 1968–1984.e20

Wu X, Ge H, Lemon B, Vonderfecht S, Weiszmann J, Hecht R, Gupte J, Hager T, Wang Z, Lindberg R, et al (2010) FGF19induced hepatocyte proliferation is mediated through FGFR4 activation. Journal of Biological Chemistry 285: 5165–5170

Ye Y & Godzik A (2003) Flexible structure alignment by chaining aligned fragment pairs allowing twists. Bioinformatics 19: ii246–ii255

Zhang G, Budker V & Wolff JA (1999) High levels of foreign gene expression in hepatocytes after tail vein injections of naked plasmid DNA. Hum Gene Ther 10: 1735–7

Zhou M, Learned RM, Rossi SJ, DePaoli AM, Tian H & Ling L (2017a) Engineered FGF19 eliminates bile acid toxicity and lipotoxicity leading to resolution of steatohepatitis and fibrosis in mice. Hepatol Commun 1: 1024–1042

Zhou M, Luo J, Chen M, Yang H, Learned RM, DePaoli AM, Tian H & Ling L (2017b) Mouse species-specific control of hepatocarcinogenesis and metabolism by FGF19/FGF15. J Hepatol 66: 1182–1192

Zhou M, Wang X, Phung V, Lindhout DA, Mondal K, Hsu JY, Yang H, Humphrey M, Ding X, Arora T, et al (2014a) Separating tumorigenicity from bile acid regulatory activity for endocrine hormone FGF19. Cancer Res 74: 3306–3316

Zhou M, Wang X, Phung V, Lindhout DA, Mondal K, Hsu JY, Yang H, Humphrey M, Ding X, Arora T, et al (2014b) Separating tumorigenicity from bile acid regulatory activity for endocrine hormone FGF19. Cancer Res 74: 3306–3316

Zhou M, Yang H, Learned RM, Tian H & Ling L (2017c) Non-cell-autonomous activation of IL-6/STAT3 signaling mediates FGF19-driven hepatocarcinogenesis. Nat Commun 8: 1–16

Zucman-Rossi J, Villanueva A, Nault JC & Llovet JM (2015) Genetic Landscape and Biomarkers of Hepatocellular Carcinoma. Gastroenterology 149: 1226–1239

