## Supplementary figure 1 for "FGF19 and its analog Aldafermin cooperate with MYC to induce aggressive hepatocarcinogenesis"

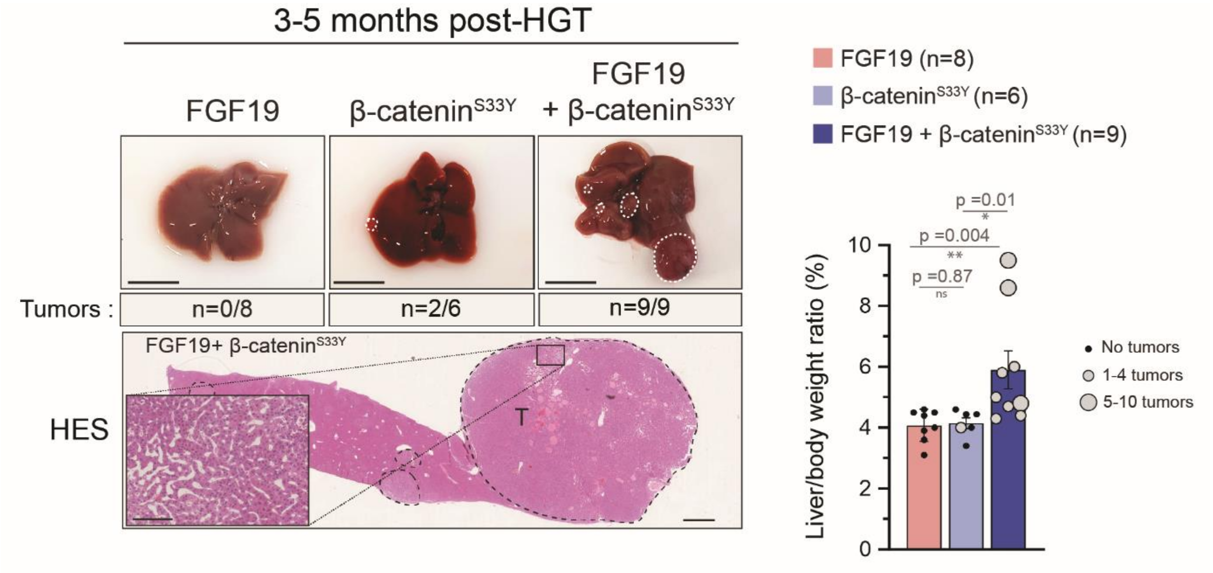


**Supplementary figure 1: FGF19 cooperates with β-cateninS33Y to induce hepatic carcinogenesis.** Representative liver and HES pictures of mice following hydrodynamic gene transfer with either FGF19, β-catenin$^{S33Y}$ or both. Liver/body weight ratio for each mice, with tumor-dependent dot sizes. Scale bar for zoom: 200µm. Scale bar for large view: 1cm. Mann-Whitney test statistical significance is indicated. Abbreviations: HES: Hematoxylin & Eosin & Saffron.
